# Non-ablative disease-modifying effects of magnetic resonance-guided focused ultrasound in neuromelanin-producing parkinsonian rodents

**DOI:** 10.1101/2023.08.08.552410

**Authors:** Joan Compte, Marion Tible, Thais Cuadros, Jordi Romero-Gimenez, Ariadna Laguna, Jean-François Aubry, Erik Dumont, Charlotte Constans, Thomas Tiennot, Mathieu D. Santin, Stephane Lehericy, Miquel Vila

## Abstract

Age-dependent accumulation of the brain pigment neuromelanin has been implicated in the pathogenesis of Parkinson’s disease (PD). In humans, intracellular and extracellular neuromelanin levels are increased in PD postmortem brains and boosting neuromelanin production in rodents compromises neuronal function and viability and triggers a PD-like phenotype. Focused ultrasound has been shown to reduce ultraviolet light-induced skin hyperpigmentation in guinea pig and to remove brain extracellular β-amyloid plaques in Alzheimer’s mouse models. Here we show that repeated application of transcranial focused ultrasound (tFUS) is able to decrease intracellular and extracellular neuromelanin levels in neuromelanin-producing parkinsonian rats, compared to sham-treated animals, without the need for any additional therapeutic agent or intervention. Reduced neuromelanin levels in tFUS-treated animals were associated with decreased Lewy-like pathology, preserved dopaminergic phenotype, attenuated nigrostriatal degeneration, reduced glial activation, and long-term recovery of motor function. Our findings indicate that tFUS treatment applied at prodromal/early disease stages provides by itself extended structural and functional preservation of the nigrostriatal pathway in neuromelanin-producing parkinsonian rats without causing overt neuronal damage. This FDA-approved technology should thus be explored further as a noninvasive method with neuroprotective potential in PD and to maintain neuromelanin to levels below its pathogenic threshold within the aging population.

**One Sentence Summary:** Transcranial focused ultrasound reduces age-dependent neuromelanin accumulation and provides therapeutic benefit in parkinsonian rats

## INTRODUCTION

Humans accumulate with age the brain pigment neuromelanin (NM). In contrast to the widespread distribution of other brain pigments such as lipofuscin, NM is restricted to catecholamine-producing neurons, as this pigment derives from the oxidation of catechols linked to catecholamine metabolism and neurotransmission^1^. Because neurons lack the capacity to degrade or eliminate this pigment, intracellular NM progressively builds up with age until occupying most of the neuronal cytoplasm^2, 3^. Importantly, neurons reaching the highest levels of NM preferentially degenerate in Parkinson’s disease (PD), in particular dopaminergic (DA) neurons of the substantia nigra (SN), the loss of which leads to the classical motor symptoms of the disease^4^. While NM does not appear spontaneously in most animals, including rodents, we recently developed the first rodent model of human-like NM production by viral vector-mediated expression of melanin-producing enzyme tyrosinase (AAV-TYR) in rat SN^5^. This has revealed that NM can trigger PD pathology when accumulated above a specific threshold, including motor deficits, Lewy body (LB)-like pathology, nigrostriatal neurodegeneration and neuroinflammatory changes^5^. Relevant to humans, intracellular NM levels reach this pathogenic threshold in PD patients and pre-PD subjects^5^. These results suggest that strategies to maintain or reduce NM levels below this threshold could potentially provide therapeutic benefit in PD. Indeed, we previously reported that reduction of intracellular NM levels with gene therapy in NM-producing animals, either by boosting NM cytosolic clearance with autophagy activator TFEB^5^ or by decreasing NM production with VMAT2-mediated enhancement of dopamine vesicular encapsulation^6^, resulted in a major attenuation of the PD phenotype in these animals, both at the behavioral and neuropathological level. However, whether NM levels could be therapeutically modulated in vivo in a noninvasive translational manner remains unexplored.

Magnetic resonance (MR)-guided transcranial focused ultrasound (tFUS) is a cutting-edge medical technology that offers the ability to non-invasively and precisely intervene in specific brain circuits through the intact skull^7^. The actions of tFUS in the brain take many forms, ranging from transient blood-brain barrier opening and neuromodulation to permanent thermoablation. In the context of PD, tFUS has arisen as a promising incisionless alternative to surgical procedures. Indeed, FDA-approved neurological indications for MR-guided tFUS include thalamotomy for tremor-dominant PD^7^. In addition, tFUS subthalamotomy in one hemisphere has been shown to improve motor features in selected PD patients with asymmetric signs^8^.

Relevant to the present work, tFUS has been shown to decrease ultraviolet light-induced hyperpigmentation in experimental animal models by mechanically eliminating melanin and pigmented debris from the epidermis and upper dermis^9^. Furthermore, tFUS has also been successfully applied to remove extracellular depositions, such as β-amyloid plaques in Alzheimer’s mouse models, by stimulating microglia-mediated clearance^10^. Based on these observations, here we assessed whether tFUS treatment might be able to decrease pathological intracellular NM levels and extracellular NM debris released from dying neurons in humanized NM-producing parkinsonian rats, as a potential noninvasive strategy to therapeutically modulate NM levels in vivo.

## RESULTS

### Early-stage MRI-guided tFUS application in NM-producing parkinsonian rats

To determine potential direct effects of tFUS treatment on NM accumulation and PD pathology we used humanized NM-producing parkinsonian rats. These animals recapitulate several features of PD including Lewy body-like inclusion formation, degeneration of nigrostriatal melanized neurons, extracellular release of NM from dying neurons, nigral inflammatory changes and motor deficits^5^. This model is based on the unilateral adeno-associated viral vector-mediated expression of melanin-producing enzyme Tyr (AAV-TYR) in the rat SN. By 1 month (m) post-AAV these animals exhibit intracellular NM levels equivalent to aged human SN^5^. At 1-2 m post-AAV, intracellular NM reaches levels equivalent to those observed in PD patients, and these animals start exhibiting a PD-like phenotype that becomes fully established by 4 m post-AAV injection^5^. Based on this temporal dynamics, we applied tFUS treatment between 1 and 2 m post-AAV, corresponding to early PD stages, and assessed the behavioral and histological effects at 4 m post-AAV (Fig. 1). Control animals were injected with an empty viral vector (AAV-EV). Both groups of animals (AAV-TYR and AAV-EV) were divided into two experimental groups for either tFUS or sham treatment (n=6 animals per treatment and group). By taking advantage of the presence of NM in AAV-TYR-injected rats, NM-sensitive high-resolution T1-weighted magnetic resonance imaging (NM-MRI) was performed to guide the application of tFUS into the SN ipsilateral to AAV injections. In all experimental groups, MRI-guided tFUS/sham treatments were performed once a week for 3 consecutive weeks. The power of the tFUS ultrasound beam was initially set at 5% corresponding to a Peak Negative Pressure of 1,05 MPa in water according to the calibration, with a duration of 3 ms every 97 ms for 600 repetitions (10 minutes), corresponding to I_spta_= 1.1W/cm^2^. The parameters were chosen to limit the thermal effect and were within the range of parameters previously used^11^, in which the maximum thermal rise was estimated to 0.12°C. At 4 m post-AAV injections, corresponding to ∼2.5 m after the last tFUS application, animals were assessed for contralateral paw akinesia (cylinder test) and processed for neuropathological examination. Overall, MRI-guided tFUS application in the SN of NM-producing rats proved feasible, reproducible, localized, and could thus be used to further investigate the influence of this treatment on the PD-like phenotype in this animal model.

**Figure 1.**
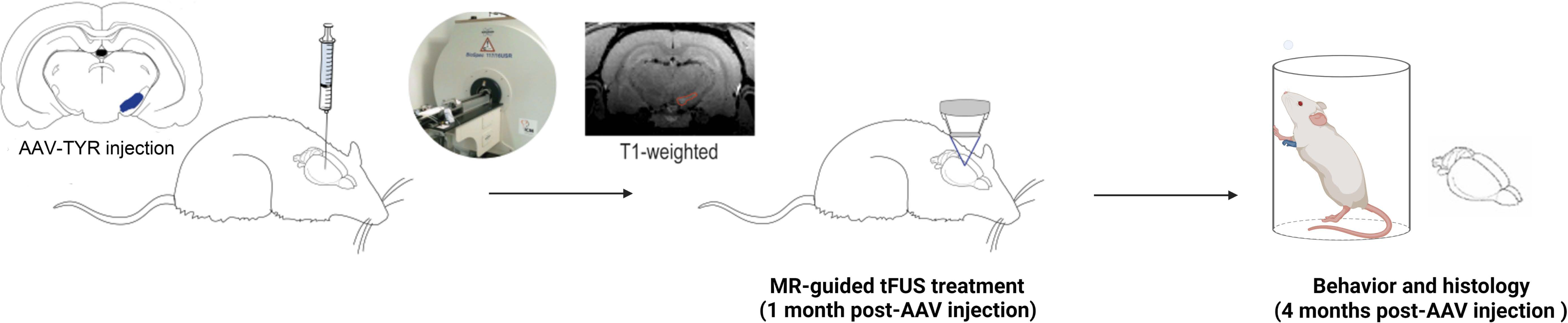
Experimental design for tFUS treatment in parkinsonian NM-producing rats using a rodent-adjustable tailor-made tFUS equipment.

### TFUS reduces intracellular NM levels and extracellular NM debris in NM-producing rats

We first assessed whether tFUS focused at the melanized SN from NM-producing rats would decrease the age-dependent accumulation of nigral NM occurring in these animals. This was determined by measuring both intracellular NM levels and extracellular NM debris released from dying neurons in the SN of NM-producing rats at 4 m post-AAV-TYR injections. At this stage, these animals have reached pathological NM levels linked to conspicuous nigrostriatal degeneration^5^. Compared to sham-treated animals, tFUS treatment was associated with a reduction of intracellular NM levels, as assessed by optical densitometry (Fig. 2A and Supplementary Fig. 1). Importantly, tFUS treatment was able to maintain intracellular NM levels below the pathogenic threshold above which we previously reported overt established neurodegeneration in these animals^5^ (Fig. 2A and Supplementary Fig. 1). Reductions in intracellular NM levels were accompanied by a decreased surface of cytosolic neuronal area occupied by NM (Fig. 2B and Supplementary Fig. 1). In tFUS-treated rats, NM pigment occupied less than 50% of the total cytosolic area while in sham-treated animals more than half of the neuronal cytosol was filled with NM (Fig. 2B and Supplementary Fig. 1). The reduction in intracellular NM levels by tFUS was accompanied by concomitant decreases in extracellular NM debris (Fig. 2B and Supplementary Fig. 1), which are abundantly observed in aged and PD postmortem human brains associated with microglial/macrophage activation^12^. In particular, tFUS treatment reduced both the number and the size of extracellular NM clusters (Fig. 2B and Supplementary Fig. 1). Overall, these results indicate that tFUS treatment is able to lessen the extent of NM buildup occurring with age, both at intracellular and extracellular level.

**Figure 2.**
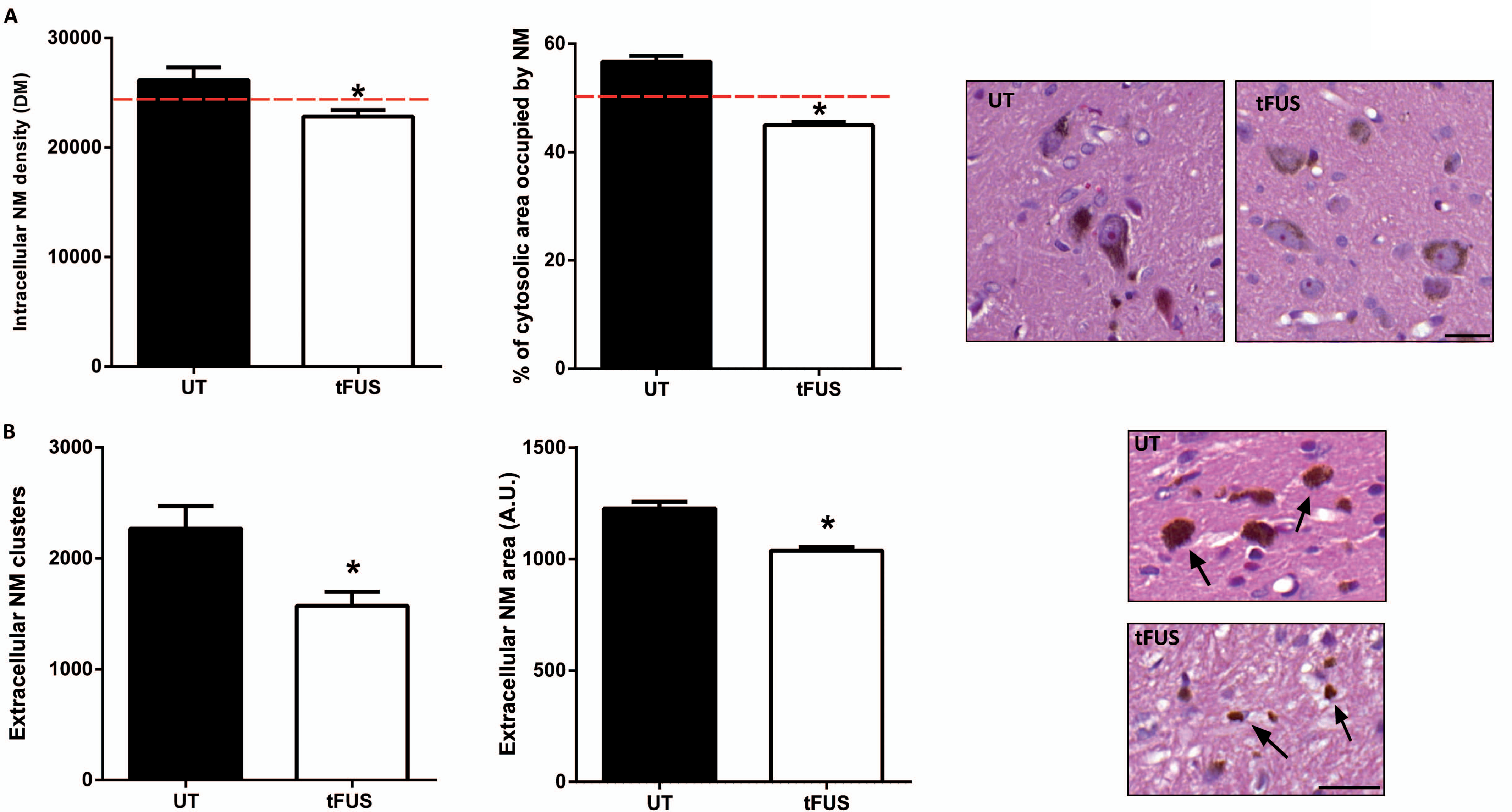
TFUS reduces intracellular and extracellular NM accumulation in NM-producing rats. **(A)** Quantification of intracellular NM levels by optical densitometry (*left*) and percentage of neuronal cytosolic area occupied by NM (*right*) in ipsilateral SN DA neurons of AAV-TYR-injected rats at 4 m post-injection, either sham-treated (UT, untreated) or treated with tFUS. Dashed red lines represent the threshold of density measurement (DM) above which we previously reported overt established neurodegeneration in these animals ^5^ (i.e. 24,000 DM, *left*) or indicate a 50% occupancy of the cytosol by NM (*right*). *Micrographs*, representative images of 5 µm-thick H&E-stained SN sections from these animals (unstained NM in brown). (**B)** Quantification of the number (*left*) and size (*right*) of extracellular NM debris in ipsilateral SN of AAV-TYR-injected rats at 4 m post injection, either sham-treated (UT) or treated with tFUS. *Micrographs*, representative images of 5 µm-thick H&E-stained SN sections from these animals (arrows, unstained extracellular NM debris, in brown). In all panels, \**p*<0.05 vs UT, Mann-Whitney test; values are mean ± SEM (individual values per group are shown in Supplementary Fig. 1). In **A**, N=206 neurons from 6 UT animals and 822 neurons from 6 tFUS-treated animals. In **B** (*left*) N=6 UT and 6 tFUS-treated animals; (*right*), N=1002 NM clusters per group, from 6 UT animals and 6 tFUS-treated animals. Scale bars: 20 µm.

### Reduced extracellular NM debris by tFUS is associated with attenuated PD-like neuroinflammation

Extracellular NM debris released from dying neurons, as observed in NM-producing parkinsonian rats, is a common PD feature often associated with activated microglia, which is indicative of an active, ongoing neurodegenerative process (i.e., neuronophagia). Microglial cells are indeed considered the main players in the recognition, engulfment and clearance of extracellular NM^12^. In agreement with this, extracellular NM in NM-producing rats was associated with a robustly increased number of Iba1-positive microglial cells with a phagocytic/reactive morphology (i.e. large amoeboid de-ramified) containing or surrounding NM debris (Fig. 3A). In these animals, tFUS-induced attenuation of extracellular NM was associated with a largely diminished microglial activation (Fig. 3A). Similarly, tFUS treatment also reduced the recruitment of CD68-positive cells, corresponding to tissue-resident or blood-borne macrophages with phagocytic activity that are found in close association with extracellular NM in NM-producing animals (Fig. 3B), as it occurs in PD brains^5, 12^. Additional inflammatory changes observed in NM-producing rats included increased astrocyte reactivity widely distributed within the SN, which was also markedly attenuated by tFUS treatment (Fig. 3C). In contrast, no inflammatory traces were observed in the SN of tFUS-treated EV control animals, indicating that tFUS *per se* did not induce brain inflammation (Fig. 3A-C). Taken together, these results indicate that, by reducing extracellular NM accumulation, tFUS treatment is able to attenuate the overall PD-like inflammatory response linked to presence of extracellular NM debris.

**Figure 3.**
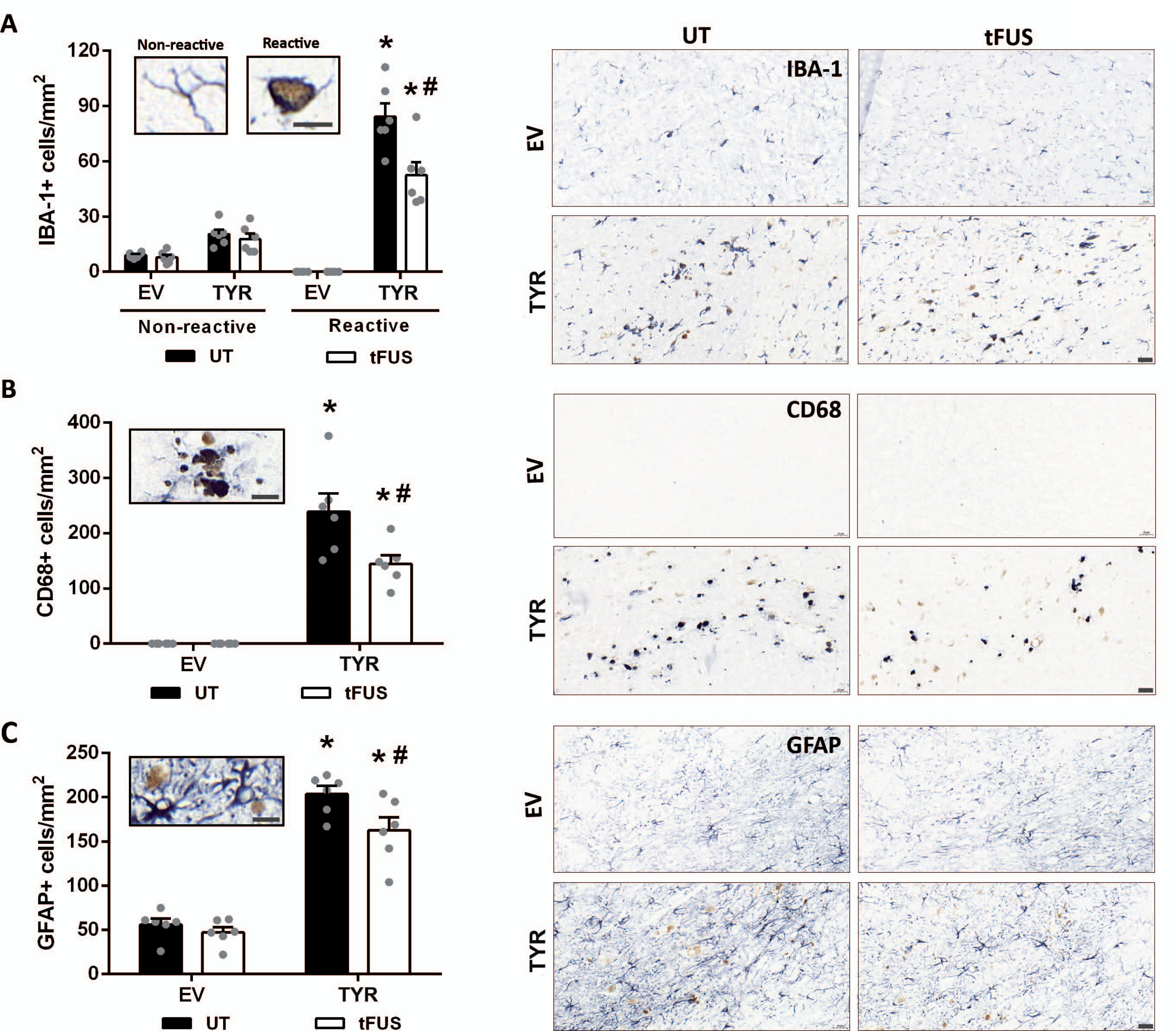
TFUS diminishes PD-like inflammatory changes in NM-producing rats. Quantification of: **(A)** Iba-1-positive microglial cells with a non-reactive (ramified) and phagocytic/reactive (large amoeboid de-ramified) morphologies, **(B)** CD68-positive macrophages, and **(C)** GFAP-positive astrocytes in the ipsilateral SN of AAV-TYR-injected melanized rats and AAV-EV-injected non-melanized controls at 4 m post-injection, either sham-treated (UT) or treated with tFUS. *Micrographs*, representative images of 5 µm-thick immunostained SN sections with Iba-1 **(A)**, CD68 (**B)** and GFAP **(C)** in blue and unstained NM in brown. In all panels, values are mean ± SEM and N=6 animals per experimental group. In **A**, \**p*<0.05 vs respective non-reactive state; ^#^*p*<0.05 vs UT reactive state; Two-way ANOVA with Tukey’s post-hoc test. In **B** & **C**, \**p*<0.05 vs respective EV group; ^#^*p*<0.05 vs UT AAV-TYR-injected animals; Two-way ANOVA with Tukey’s post-hoc test. In all panels, N=6 animals per group. Scale bars: 20 µm & 10 µm (insets).

### TFUS attenuates NM-linked nigral neurodegeneration

To determine whether tFUS-induced reductions in NM levels and neuroinflammation impacts nigral neuronal viability, we assessed DA neuronal integrity in tFUS-treated parkinsonian rats. First, we verified that tFUS treatment by itself was not deleterious to neurons, as indicated by a lack of effect of tFUS on the morphology and number of DA nigral neurons in non-melanized control rats (Fig. 4A). In NM-producing parkinsonian animals, tFUS significantly attenuated the loss of TH-positive SN neurons that took place by 4 m post-AAV-TYR injection in these animals (Fig. 4A). Part of this effect could be attributed to an attenuation by tFUS of the phenotypic loss of TH expression that occurs within NM-laden neurons at early stages of neurodegeneration (Fig. 4B), as seen in PD brains^13^. To distinguish between the effects of tFUS on TH expression and on actual cell death, we also assessed the total number of SN DA neurons independently of the TH phenotype (Fig. 4C). This analysis revealed a structural preservation of total DA neurons by tFUS in NM-producing parkinsonian rats compared to untreated animals (Fig. 4C). In addition, preserved neurons in tFUS-treated rats exhibited fewer LB-like cytoplasmic inclusions (Fig. 4D) and less cellular atrophy/shrinkage (Fig. 4E) than those from untreated animals, which, together with the attenuated TH downregulation reported above, suggests a functional preservation of these neurons following tFUS treatment. Overall, these results indicate that tFUS mitigates PD-like neuropathology and neurodegeneration linked to age-dependent NM accumulation.

**Figure 4.**
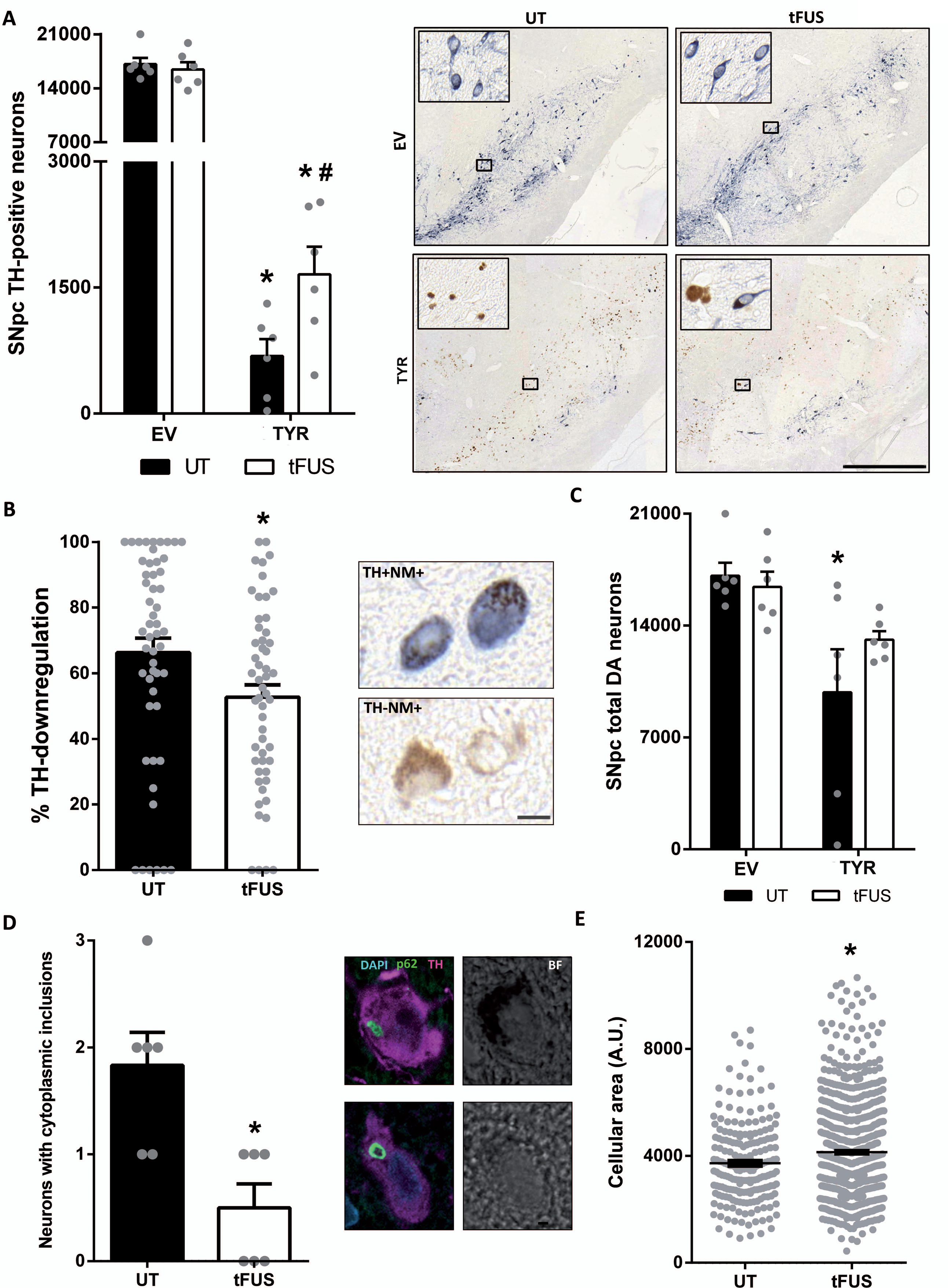
TFUS attenuates PD-like neuropathology in NM-producing parkinsonian rats. **(A)** Stereological cell counts of TH-positive neurons in the ipsilateral SN of AAV-TYR-injected rats and AAV-EV-injected non-melanized controls at 4 m post-AAV injections, either sham-treated (UT) or treated with tFUS. *Micrographs*, representative images of 5 µm-thick TH-immunostained ipsilateral SN sections, with TH in blue and unstained NM in brown; Scale bar: 500 µm. **(B)** Percentage of pigmented nigral neurons with downregulated TH expression (i.e. TH^-^NM^+^ neurons vs the total number of NM^+^ neurons) in AAV-TYR-injected rats at 4 m post-AAV injections. *Micrographs*, representative images of TH^+^NM^+^ and TH^-^NM^+^ neurons in 5 µm-thick TH-immunostained ipsilateral SN sections, with TH in blue and unstained NM in brown. **(C)** Stereological cell counts of total number of SN DA neurons (including TH^+^NM^+^, TH^+^NM^-^ and TH^-^NM^+^) in the ipsilateral SN of AAV-TYR-injected rats and AAV-EV-injected controls at 4 m post-AAV injections, either sham-treated (UT) or treated with tFUS. **(D)** Quantification of the number of NM-laden neurons exhibiting p62-positive LB-like cytosolic inclusions in the ipsilateral SN of AAV-TYR-injected rats at 4 m post-AAV injections, either sham-treated (UT) or treated with tFUS. *Micrographs*, representative confocal images of a melanized TH^+^ neuron with a LB-like inclusion immunopositive for p62 (green; TH in pink) in the SN of an AAV-TYR-injected rat; BF, brightfield; Scale bar: 10 µm. **(E)** Quantification of the cellular area of NM-containing neurons in the ipsilateral SN of AAV-TYR-injected rats at 4 m post-AAV injections, either sham-treated (UT) or treated with tFUS. In **A-D** values are mean ± SEM. In **A**, \**p*<0.05 vs respective EV animals; ^#^*p*<0.05 vs UT AAV-TYR-injected animals. In **B**, **D** & **E**, \**p*<0.05 vs UT animals. In **C**, \**p*<0.05 vs UT EV-injected animals. **A** & **C**, Two-way ANOVA with Tukey’s post-hoc test; **B**, **D** & **E**, Mann-Whitney test. **A-D**, N=6 animals per group. In **E**, N=206 neurons from 6 untreated animals and 822 neurons from 6 tFUS-treated animals.

### TFUS reduces motor impairment in NM-producing parkinsonian rats

We next determined whether the neuroprotective effects exerted by tFUS on nigral DA neurons were accompanied by an attenuation of motor impairment in NM-producing parkinsonian rats. Consistent with preservation of nigral cell bodies, tFUS treatment was associated with a reduction in striatal DA fiber loss in these animals (Fig. 5A). This protective effect was mostly limited to the dorsolateral striatum (Supplementary Fig. 2), which receives its DA innervation exclusively from the SN and is considered to be the equivalent of the putamen in humans^14^. This striatal subregion presents the most profound DA depletion in patients with PD^15^ and appears to be critical for motor function ^16^. Accordingly, tFUS treatment markedly attenuated contralateral forepaw hypokinesia in NM-producing rats up to 4 m post-AAV-TYR injection, as assessed by the cylinder test (Fig. 5B). The level of motor function preservation for each animal significantly correlated with the level of preservation of striatal DA fibers (Fig. 5C). Overall, these results indicate that tFUS treatment at prodromal/early (pre-symptomatic) PD stages provides a long-term functional preservation of the nigrostriatal circuit in parkinsonian rats.

**Figure 5.**
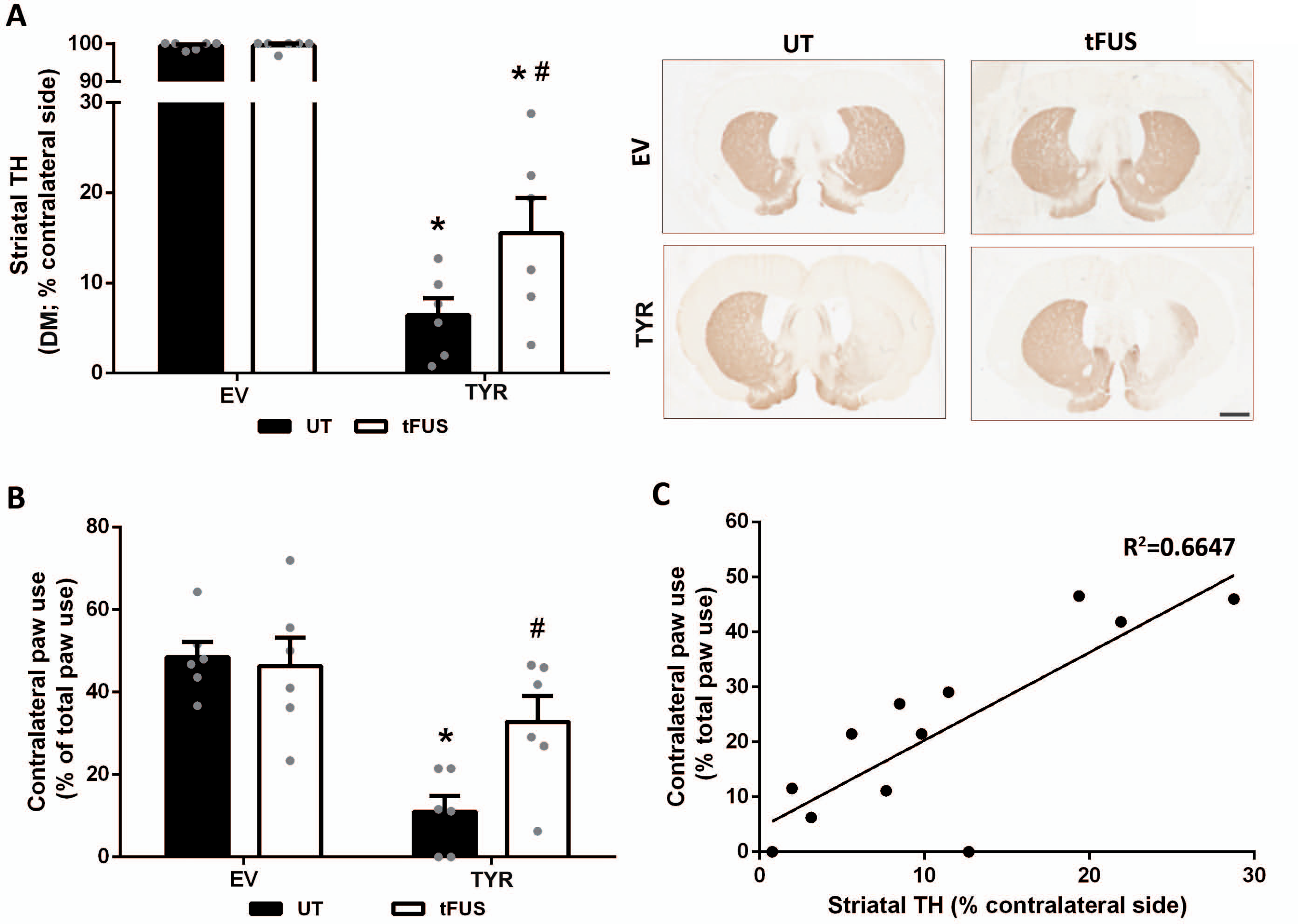
TFUS reduces nigrostriatal denervation and motor impairment in NM-producing parkinsonian rats. **(A)** Optical densitometry of striatal TH-positive fibers in AAV-TYR-injected rats and AAV-EV-injected non-melanized controls at 4 m post-AAV injections, either sham-treated (UT) or treated with tFUS. *Micrographs*, representative images of 5 µm-thick TH-immunostained striatal sections (TH in brown; right, ipsilateral hemisphere). Scale bar: 1 mm. \**p*<0.05 vs respective EV animals; ^#^*p*<0.05 vs UT AAV-TYR-injected animals; Two-way ANOVA with Tukey’s post-hoc test. **(B)** Contralateral forepaw use in AAV-TYR-injected rats and AAV-EV-injected non-melanized controls, either sham-treated (UT) or treated with tFUS at 4 m post-AAV injections. \**p*<0.05 vs respective EV; ^#^*p*<0.05 vs UT AAV-TYR-injected animals; Two-way ANOVA with Tukey’s post-hoc test. **(C)** Pearson correlation analysis between striatal TH DM (% of contralateral side) and contralateral paw use (Pearson *r*=0.815; *p*=0.001). In all panels, values are mean ± SEM and N=6 animals per experimental group.

## DISCUSSION

Using rodent-adjustable tailor-made tFUS equipment, we found that the application of intermediate-intensity tFUS was able to reduce NM levels below their pathogenic threshold and provide extended therapeutic benefit in a humanized NM-producing PD rat model (Fig. 6). Remarkably, the beneficial effects of tFUS were observed in the absence of any additional therapeutic agent or intervention, thus being compatible with a direct mechanical/thermal fragmentation of NM by tFUS, as it has been reported in the skin of hyperpigmented animal models^9^. Supporting this concept, tFUS-treated animals exhibited fragmented, smaller extracellular NM debris, which are more likely to be phagocytosed by microglia, and a decreased surface of cytosolic neuronal area occupied by intracellular NM, compared to sham-treated animals. Yet, the tFUS intensity used in our study was within the range used for either tissue ablation or neuromodulation, thus a potential contribution of such mechanisms could also be considered. However, our tFUS treatment not only produced no tissue damage or inflammation in control rats but also reduced cell death in parkinsonian animals, in contrast to the coagulative necrotic lesions induced by high-intensity tFUS for current PD applications (i.e. thalamotomy, subthalamotomy, pallidotomy). In addition, while tFUS can both inhibit and enhance neural activity in superficial and deep brain regions^7^, tFUS neuromodulatory effects are usually transient, lasting for somewhere between half an hour and a day post-treatment^17–, 19^, as opposed to the extended effects reported here up to at least 2.2 months after the last tFUS application. Therefore, while neither neuromodulatory nor ablative mechanisms can be completely ruled out, it seems that they do not play a major role in our study.

**Figure 6.**
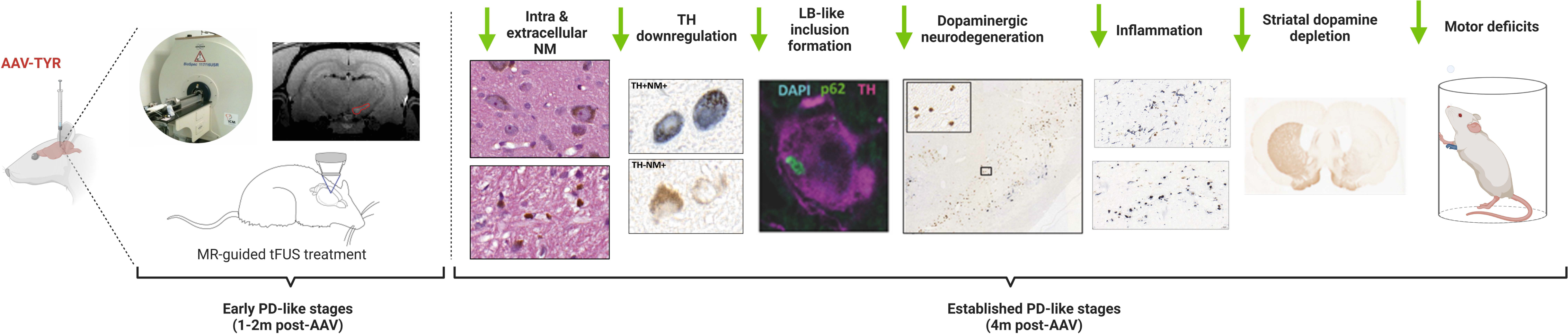
Summary schematics of the beneficial effects of tFUS treatment in parkinsonian NM-producing rats.

Instead, our results point to a non-ablative disease-modifying effect of tFUS involving a therapeutic reduction of NM levels. Supporting a deleterious role of excessive NM accumulation, we previously found that the continuous intracellular build-up of NM until occupying most of the neuronal cytoplasm ultimately leads to a general failure of cellular proteostasis coursing with lysosomal/autophagy dysfunction, ubiquitin-proteasome impairment, alpha-synuclein oligomerization, reduced mitochondrial respiration, increased production of reactive oxygen species, and cell death^5, 20^. Therefore, while it has been postulated that NM synthesis may have a beneficial role by removing potentially toxic dopamine-derived oxidative species from the cytosol, its accumulation above a pathogenic threshold may interfere with normal cell function and trigger PD-like pathology, regardless of whether the initial synthesis of NM is a putative protective process. Supporting this concept, we have previously shown that reducing intracellular NM levels below this pathogenic threshold with gene therapy markedly attenuated the PD phenotype of AAV-TYR-injected NM-producing rats^5, 6^. In the present study, while the noninvasive reduction of NM levels by tFUS was not apparently extensive, it was however sufficient to maintain NM below its pathogenic threshold of accumulation and thus may account for the attenuation of the PD phenotype in these animals. It is also important to note that NM levels in tFUS-treated animals were assessed 2.2 months after the last tFUS application, during which time NM continued to be steadily synthesized and accumulated. If NM levels were to be assessed shortly following the last tFUS application, the effects of tFUS on reducing NM levels would thus probably prove larger.

In any event, the extent of NM reduction by ultrasound could potentially be enhanced by combining tFUS with the systemic injection of microbubbles that expand and contract when activated with ultrasound to temporarily open the blood-brain barrier (BBB). BBB opening is currently one of the most extensively investigated tFUS applications, with safety and efficacy having been validated in several preclinical studies, from rodents to non-human primates in both health and disease^21^. For instance, tFUS-induced BBB opening has been successfully applied to therapeutically remove extracellular β-amyloid plaques in mouse models of Alzheimer’s disease by stimulating microglia-mediated clearance of the plaques^10^. A similar approach could thus be envisaged to help remove even further PD-linked extracellular NM debris from the brain. In addition, tFUS-induced BBB opening can help deliver therapeutic agents into the brain, including drugs, neurotrophic factors or viral vectors for gene therapy^7^. Accordingly, this approach could be used to deliver viral vectors designed to further reduce intracellular NM levels such as autophagy activator TFEB, which restores proteostasis and boosts NM cytosolic clearance^5^, or VMAT2, which enhances dopamine vesicular encapsulation and reduces NM production^6^. Importantly, tFUS-induced BBB opening has been recently proved to be feasible, safe, reversible, and can be performed repeatedly in humans^22^, so these interventions could be readily translated into the clinics.

There are, however, some limitations in the present study that should be addressed before its potential translation to humans. For instance, because tFUS treatment was applied at a prodromal/early disease stage in NM-producing animals, it remains to be determined whether in established PD patients, in which intracellular NM has already reached overt pathological levels^5^, this approach would be able to halt or delay disease progression. Also, while we have shown a beneficial effect of tFUS up to 2 months post-treatment, it is not known how much longer this beneficial effect could potentially last and whether repeated tFUS exposures would be required. In addition, PD is known to involve the accumulation of pathological alpha-synuclein in multiple brain regions beyond those that accumulate NM, thus raising the question as to whether by solely targeting melanized PD-vulnerable regions (including the SN, locus coeruleus or dorsal vagal complex) would be sufficient to have a beneficial effect in PD. In this context, alpha-synuclein has been shown to redistribute to the NM pigment in early stages of PD and become entrapped within NM granules, which may predispose melanized neurons to precipitate alpha-synuclein around pigment-associated lipids under oxidative conditions, such as those linked to NM formation^23–25^. Along this line, enhanced oxidative stress has been reported to experimentally increase in vivo neuron-to-neuron alpha-synuclein spreading from dorsal vagal complex to more rostral brain regions^26^. We had previously found that ablating alpha-synuclein in AAV-TYR-injected rodents was not sufficient to provide therapeutic benefit while reducing intracellular NM levels in these animals resulted in a major attenuation of the PD phenotype^5, 6^. Along this line, recent trials using anti-alpha-synuclein monoclonal antibodies failed to provide significant clinical benefit^27, 28^. While several reasons could explain this lack of clinical efficacy^29^, alpha-synuclein accumulation in fact correlates poorly with clinical symptoms and neurodegeneration in PD^30^. In this context, age-dependent NM accumulation, not only in the SN but also in other melanized PD-vulnerable systems, could represent an alternative, disease-relevant target with therapeutic potential.

Despite the remaining open questions, the results presented here demonstrate the potential of tFUS as a non-ablative disease-modifying treatment for early PD stages involving the therapeutic reduction of NM levels. Because all humans accumulate NM with age, such an intervention could also be potentially envisaged within the aging population to prevent NM from reaching pathological levels. Indeed, NM-filled neurons from apparently healthy aged individuals exhibit early signs of neuronal dysfunction/degeneration even in the absence of overt PD^12, 13, 31, 32^, thus implying that all humans would potentially develop PD if they were to live long enough to reach pathogenic NM levels. In fact, PD is the fastest growing neurological disorder in the world, with the number of PD patients expected to double to over 12 million worldwide by 2040^33^. In this context, interventions to prevent, halt or delay disease progression, which are currently lacking, should have a major global impact.

## MATERIALS AND METHODS

### Animals

Adult male Sprague–Dawley rats (Charles River), 225–250 g at the time of surgery, were housed two to three per cage with ad libitum access to food and water during a 12 hours (h) light/dark cycle. All the experimental and surgical procedures were conducted in accordance with the European (Directive 2010/63/UE) and Spanish laws and regulations (Real Decreto 53/2013; Generalitat de Catalunya Decret 214/97) on the protection of animals used for experimental and other scientific purposes, and approved by the Vall d’Hebron Research Institute (VHIR) Ethical Experimentation Committee. A total of 24 rats were randomly distributed into the different experimental groups (N=6 animals per experimental group, as follows: N=6/24 EV:UT, N=6/24 EV:tFUS, N=6/24 TYR:UT, N=6/24 TYR:tFUS). Control and experimental groups were processed at once to minimize bias.

### Stereotaxic infusion of viral vectors

Recombinant AAV vector serotype 2/1 expressing the human tyrosinase cDNA driven by the cytomegalovirus (CMV) promoter (AAV-TYR; concentration 2.6 × 10^13^ gc/mL) and the corresponding control empty vector (AAV-EV; concentration 2.5 × 10^13^ gc/mL) were produced at the Viral Vector Production Unit of the Autonomous University of Barcelona (UPV-UAB, Spain) using a TYR plasmid kindly provided by Dr. Takafumi Hasegawa (Department of Neurology, Tohoku University School of Medicine, Sendai, Japan). Surgical procedures were performed with the animals placed under general anesthesia using isoflurane (5% for the induction phase and 2% for the maintenance phase) (Baxter). Vector solutions were injected using a 10 μL Hamilton syringe fitted with a glass capillary (Hamilton model Cat#701). Animals received 2 μL of AAV-TYR or 2 µl AAV-EV. Infusion was performed at a rate of 0.4 μL/min and the needle was left in place for an additional 4 min period before it was slowly retracted. Injection was carried out unilaterally on the right side of the brain at the following coordinates (flat skull position), right above the SN: antero-posterior: −5.29 mm; medio-lateral: −2 mm; dorso-ventral: - 7.6 mm below dural surface, calculated relative to bregma according to the stereotaxic atlas of Paxinos and Watson^34^.

### Transduction efficiency

AAV-TYR transduction efficiency was determined at 2–4 weeks post-AAV injection (n = 6–8 rats) either by: (i) assessing TYR expression within SNpc tyrosine hydroxylase (TH)-positive cells by double immunofluorescence with antibodies against TH and TYR in six coronal midbrain sections through the entire SNpc, using the Cell Counter plugin on ImageJ software (Rasband, W.S., ImageJ, U. S. National Institutes of Health, Bethesda, Maryland, USA, 1997-2016); (ii) by counting the number of NM-positive neurons versus the total number of SNpc TH-positive neurons, as described below.

### MRI-guided tFUS

Rat weighing 300-350g at the time of the experiment were anesthetized with 4% Isoflurane and a mixture of air and oxygen (5/1) and stabilized at 1.5% Isoflurane. Their heads were closely shaved and depilated to facilitate the contact between the skin and the transducer latex membrane. A 22G catheter was positioned in the caudal vein and a bolus of 100 µL 10% heparin (Heparin Choay, 25000 UI/5ml) was delivered to avoid clot formation. Following the rat tail-first installation in the adapted MRI bed, the tFUS transducer was placed on its head with the stereotactic setup described in ref.^35^. Acoustic coupling of the transducer with the head was achieved by a degassed, deionized water-filled inflatable balloon covered with a thin layer of acoustic gel (Aquasonic 100, Parker Laboratories Inc., Fairfield, NJ, USA). TFUS calibration measurements were performed with a needle hydrophone (HNC 400, SN 1207, Onda Corp, Sunnyvale Ca, USA) connected to a numerical oscilloscope (Handyscope HS5, Tiepie Engineering, Sneek, the Netherlands). For each treatment the tFUS apparatus was positioned based on anatomical brain MRI. Three stacks of 2D slices were acquired with an 11.7T MRI (Bruker Biospec 117/16, Germany), each one with different orientation (axial, coronal and sagittal) using a 2D Gradient Echo sequence. Imaging parameters were the same for the 3 acquisitions: FOV = 51.2*51.2 mm^2^, with 20 contiguous slices of 1.5 mm thickness, matrix size = 256*256 (200 µm isotropic in plane resolution), TR/TE = 300 ms/4 ms and flip angle θ = 60°. Each scan lasted 1 min 17 s. A tFUS trajectory was then planned thanks to these sequences using the Thermoguide software (IGT, Pessac, FRANCE) and tFUS were sent thanks to a 20 mm curvature radius 7-elements 1, 5 MHz transducer (Imasonic, Voray sur l’Ognon, France). The apparatus was positioned with MR compatible motors to reach the targeted area. The power of the tFUS ultrasound beam was initially set at 5% corresponding to a Peak Negative Pressure of 1,05 MPa in water according to the calibration, with a duration of 3 ms every 97 ms for 600 repetitions (10 minutes), corresponding to I_spta_= 1.1W/cm^2^.

### Cylinder behavioral test

Rats were tested for left and right forepaw use with the cylinder test one week before surgery and four months after viral injection. The number of animals for each condition was EV:UT (*n*=6), EV:tFUS (*n*=6), TYR:UT (*n*=6), TYR:tFUS (*n*=6). For the performance of the cylinder test, rats were first allowed to habituate to the experimental room for at least 1 h before each test. Then, rats were put in a glass cylinder and the total number of left and right forepaw touches performed within 5 min was counted. Data are presented as the percentage of the contralateral paw usage of AAV-EV and AAV-TYR-injected rats with respect to their non-melanized counterparts, either UT or tFUS. Behavioral equipment was cleaned with 70% ethanol after each test session to avoid olfactory cues. All behavioral tests were performed during the light cycle by an investigator blinded to the experimental groups.

### Brain processing for histological analyses

Four months after injection of viral vectors animals were euthanized. To this purpose, animals were deeply anesthetized with sodium pentobarbital (50 mg/kg, i.p.) and then perfused through the left ventricle with saline [0.9% (wt/vol)] at room temperature (RT), followed by ice-cold formaldehyde solution 4% phosphate buffered for histology (Panreac). The brains were removed and post-fixed for 24 h in the same fixative and subsequently processed for paraffin embedding following standard procedures. Sectioning was performed with a sliding microtome (Leica, Germany) at 5-μm-thickness. Standard hematoxylin-eosin (H&E) staining was performed in 5-μm-thick paraffin-embedded SNpc section for each animal. In these sections, SNpc DA neurons were identified by the visualization of unstained NM pigment.

### Immunohistochemistry

Deparaffinized rat brain sections were quenched for 10 min in 3% H_2_O_2_-10% (vol/vol) methanol. Antigen retrieval in paraffin sections was performed with a 10 mM citric acid solution at pH 6.0 in a microwave for 20 min. Sections were rinsed 3 times in 0.1 M Tris buffered saline (TBS) between each incubation period. Blocking for 1 h with 5% (vol/vol) normal goat serum (NGS, Vector Laboratories) was followed by incubation with the appropriate primary antibody at 4°C for 48 h in 2% (vol/vol) serum. Sections were then incubated with the corresponding secondary biotinylated antibody (Vector Laboratories), visualized by incubation with avidin-biotin-peroxidase complex (Ultrasensitive and Immunopure ABC Peroxidase staining kits for the striatum and for the SNpc, respectively; Thermo Fisher Scientific) using the VectorSG Peroxidase Substrate Kit (Vector Laboratories) as a chromogen, and mounted and coverslipped with DPX mounting medium (Sigma-Aldrich). Primary antibodies and dilution factors were as follows: rabbit anti-TH (1:40000 for SNpc and 1:3500 for striatum immunochemistry; 1:1000 for immunofluorescence, Calbiochem, Cat#657012); mouse anti-GFAP (1:1000, Sigma-Aldrich, Cat#G3839), rabbit anti-Iba1 (1:1000, Wako, Cat#019-19741); mouse anti-CD68 (1:100, Serotec, Cat#MCA341R); guinea pig anti-p62 (1:1000, Progen, Cat#GP62).

### Immunofluorescence

Immunofluorescence procedure was similar to the previously reported immunohistochemistry protocol without the quenching step. Blocking was performed with 5% (vol/vol) NGS and 0.1% (vol/vol) Triton X-100 (Sigma-Aldrich) in phosphate buffered saline (PBS) solution. Corresponding primary antibodies were incubated together overnight at 4°C in 2% (vol/vol) serum. Adequate Alexa 488, 569, and/or 647-conjugated secondary antibodies (1:1000, Thermo Fisher Scientific) were then incubated simultaneously for 1 h at RT in 2% (vol/vol) serum. Nuclei were stained with Hoechst 33342 (1:2000, Thermo Fisher Scientific) in 1× PBS for 10 min. Sections were coverslipped using the Dako Cytomation Fluorescent Mounting Medium (Dako). Immunofluorescent images were taken with a LSM 980 with Airyscan 2 confocal microscope (Zeiss, Germany) and were analyzed with ZEN 3.1 software (Zeiss, Germany). The total number of p62-positive cytoplasmic inclusions was determined in six SNpc sections per animal, from two different experimental groups: UT (*n*=6) and tFUS (n=6), at two months post-AAV injections. All quantifications were performed by an investigator blinded to the experimental groups.

### Stereological cell counting

For slide visualization a Zeiss Imager.D1 microscope coupled to an AxioCam MRc camera (Zeiss, Germany) was employed. Assessment of the total number of TH-positive neurons, the number of NM-laden neurons (with or without TH), and extracellular NM aggregates in the SNpc was performed according to the fractionator principle, using the MBF Bioscience StereoInvestigator 11 (64 bits) Software (Micro Brightfield). Serial 5 µm-thick paraffin sections covering the entire SNpc were included in the counting procedure (every 17^th^ section for a total of 10-12 sections analyzed/animal). The following sampling parameters were used: (i) a fixed counting frame with a width and length of 50 μm; (ii) a sampling grid size of 115 × 70 μm and (iii) a multiplication factor of 17. The counting frames were placed randomly by the software at the intersections of the grid within the outlined structure of interest. Objects in both brain hemispheres were independently counted following the unbiased sampling rule using a 100× lens and included in the measurement when they came into focus within the dissector. A coefficient of error of <0.10 was accepted. Data for the total numbers of TH-positive neurons and NM-containing neurons are expressed as absolute numbers for each hemisphere. The total number of SNpc DA neurons was calculated by considering all TH^+^NM^+^, TH^-^NM^+^ and TH^+^NM^-^ neurons. The percentage of TH downregulation was calculated by considering the total number of TH^+^NM^+^ and the total number of TH^-^NM^+^ with respect to the total number of nigral DA neurons containing NM. All four groups (EV:UT, EV:tFUS; TYR:UT and TYR:tFUS) were analyzed at an *n*=6. All quantifications were performed by an investigator blinded to the experimental groups.

### Quantification of neuropathological parameters

The number of p62-immunopositive Lewy body-like inclusions was counted from SNpc sections fluorescently immunostained with anti-p62 and anti-TH. In order to be taken into account, p62^+^ inclusions had to co-localize within TH^+^ neurons and needed to be cytoplasmic, not nuclear. The total number of p62-positive inclusions falling into each category was counted from images covering the whole SNpc region in each section. Five to six sections per animal were quantified. Quantifications were performed in AAV-TYR:UT and AAV-TYR:tFUS at an *n*=6. All quantifications were performed by an investigator blinded to the experimental groups.

### Quantification of neuroinflammation parameters

Quantification of Iba-1, CD68 and GFAP-positive cells was performed in SNpc sections adjacent to those used for stereological cell counts. Three slides per animal were scanned using the Panoramic Midi II FL, HQ SCIENTIFIC 60x scanner (3D Histech, Hungary) and section images were acquired with CaseViewer software (3D Histech, Hungary). For quantification of Iba-1, CD68 and GFAP-positive cells, specific artificial intelligence (AI)-based algorithms were created using the Aiforia platform (Aiforia Technologies, Finland). Iba-1-positive cells were counted separately in two different groups according to their activation state: non-reactive (branched-shaped) and reactive (amoeboid-shaped). CD68 and GFAP-positive cells were counted individually. All quantifications were performed by an investigator blinded to the experimental groups.

### Intracellular and extracellular NM analysis

Intracellular NM levels were quantified in injected animals at two months post-AAV injections in 5 µm-thick paraffin-embedded H&E-stained sections covering the whole SNpc for each animal. Eight sections were analyzed per animal for each group of the AAV-TYR-injected animals: 6x TYR:tFUS and 6x TYR:EV. In these sections, SNpc dopaminergic neurons were identified by the visualization of unstained NM brown pigment. Midbrain sections were scanned using the Pannoramic Midi II FL, HQ SCIENTIFIC (3DHistech, Hungary) and section images were acquired with CaseViewer software at an objective magnification of ×63. Quantification of the intracellular density of NM pigment was achieved by means of optical densitometry using ImageJ software (NIH, USA) as previously reported^5, 36^. Briefly, the pixel brightness values for all individual NM-positive cells (excluding the nucleus) in all acquired images were measured and corrected for non-specific background staining by subtracting values obtained from the neuropil in the same images. Similarly, extracellular NM granules pigments density measured by means of Image J. In this case, density values were directly taken without the necessity of normalization. All quantifications were performed by an investigator blinded to the experimental groups.

### Optical densitometry analyses

The density of TH-positive fibers in the striatum was measured by densitometry in serial coronal sections covering the whole region (10 sections/animal). TH-immunostained 5-μm-thick paraffin-embedded sections were scanned with an Epson Perfection v750 Pro scanner and the resulting images were quantified using Sigma Scan Pro 5 software (Systat Software Inc, USA). Striatal densitometry values were corrected for non-specific background staining by subtracting densitometric values obtained from the cortex. Data are expressed as the percentage of the densitometric value of the equivalent anatomical area from the non-injected contralateral side of the same animal. All four groups (EV:UT, EV:tFUS; TYR:UT and TYR:tFUS) were analyzed at an *n*=6. All quantifications were performed by an investigator blinded to the experimental groups.

### Statistical analysis

Statistical analyses were performed with GraphPad Prism software (v8, GraphPad Software Inc, USA) using the appropriate statistical tests, as indicated in each figure legend. No statistical methods were used to pre-determine sample size but our sample size (all groups were *n*=6) were equivalent to those reported in previous similar publications^31^. Outlier values were identified by the ROUT test and excluded from the analyses when applicable. Selection of the pertinent data representation and statistical test for each experiment was determined after formally testing for normality with the Shapiro-Wilk normality test. Accordingly, differences among means or medians were analyzed either by 1- or 2-way analysis of variance (ANOVA), Kruskal–Wallis ANOVA on ranks or Mann–Whitney rank sum test as appropriate, after formally testing for normality with the Shapiro-Wilk normality test, and indicated in each figure legend. When ANOVA showed significant differences, pairwise comparisons between means were subjected to Tukey’s post-hoc testing for multiple comparisons. Values are expressed either as mean ± standard error of the mean (SEM) or presented as box plots, with minimum, maximum and median indicated, depending on the performance of parametric or non-parametric analyses, respectively. In all analyses, the null hypothesis was rejected at the 0.05 level.

## Funding

This work was funded in whole or in part by: Aligning Science Across Parkinson’s through The Michael J. Fox Foundation for Parkinson’s Research, USA (ASAP-020505 to MV); The Michael J. Fox Foundation for Parkinson’s Research, USA (MJFF-007184 and MJFF-001059 to MV); Ministry of Science and Innovation (MICINN), Spain (PID2020-116339RB-I00 to MV); EU Joint Programme Neurodegenerative Disease Research (JPND), Instituto de Salud Carlos III, EU/Spain (AC20/00121 to MV) and Agence Nationale de la Recherche (to SL); Centres of Excellence in Neurodegeneration (CoEN4016 to MV and SL); Bettencourt Schueller Foundation, Paris, France (UltraBrain project to SL and JFA); the ‘Investissements d’Avenir’ program of the National Agency for Research (references ANR-10-EQPX-15, IAIHU-06 Paris Institute of Neurosciences-IHU to JFA and SL), the Paris Institute of Translational Neuroscience (IAIHU-06 to SL); France Life Imaging (ANR-11-INBS-0006 to SL); NeurATRIS (translational research infrastructure for biotherapies in neuroscience, reference ANR-11-INBS-0011 to SL); La Caixa Bank Foundation, Spain (INPhINIT fellowship, code LCF/BQ/DI18/1166063 to JC; Junior Leader Fellowship LCF/BQ/PR19/11700005 to AL; Health Research Grant, ID 100010434 under the agreement LCF/PR/HR17/52150003 to MV).

## Author contributions

Conceptualization: MV, SL; Methodology & Investigation: JC, MT, TC, JRG, AL, JFA, ED, CC, TT, MDS, SL, MV; Funding acquisition: MV, SL; Supervision: MV, SL, MDS; Writing – original draft: JC, MV; Writing – review & editing: All Authors

## Competing interests

Authors declare that they have no competing interests.

## Data and materials availability

All data are available in the main text or the supplementary materials.

---

**Supplementary Figure 1.**
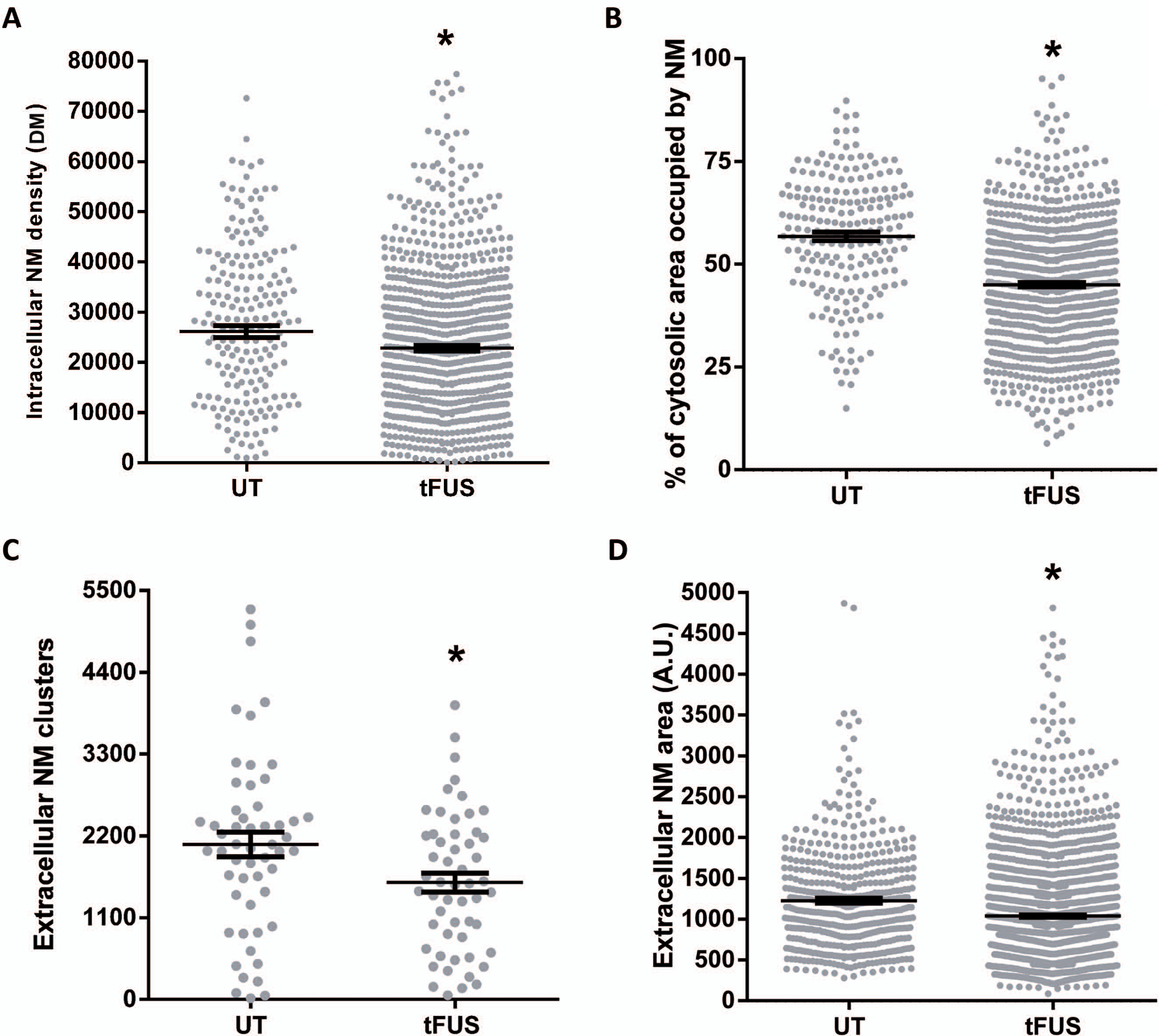
TFUS reduces intracellular and extracellular NM accumulation in NM-producing rats. Individual values per group, corresponding to the histograms shown in Fig. 2, of intracellular NM levels by optical densitometry **(A)**, percentage of neuronal cytosolic area occupied by NM **(B)**, and number **(C)** and size **(D)** of extracellular NM debris, in ipsilateral SN of AAV-TYR-injected rats at 4 m post-injection either sham-treated (UT) or treated with tFUS. In all panels, \**p*<0.05 vs UT, Mann-Whitney test. **A-B**, N=206 neurons from 6 UT animals and 822 neurons from 6 tFUS-treated animals; **C**, N=6 UT and 6 tFUS-treated animals; **D**, N=1002 NM clusters per group, from 6 UT animals and 6 tFUS-treated animals. Mean ± SEM are indicated in each panel for reference.

**Supplementary Figure 2.**
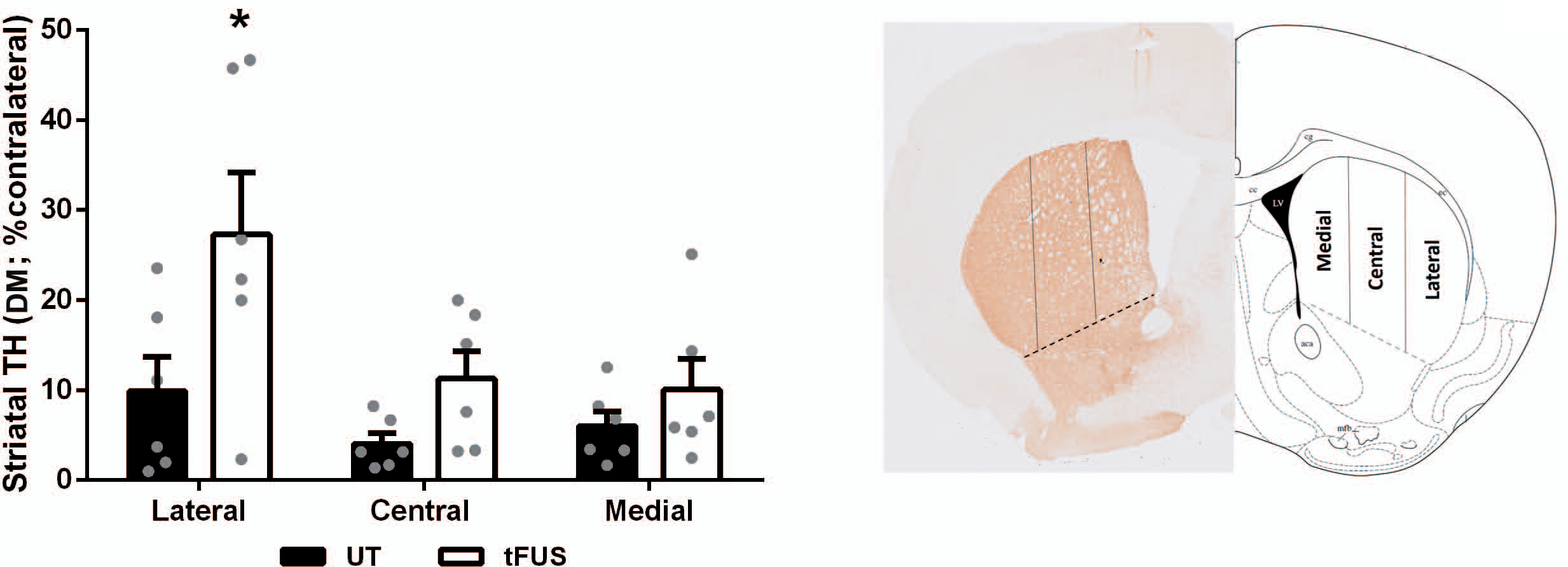
TFUS reduces dorsolateral nigrostriatal denervation in NM-producing parkinsonian rats. Optical densitometry of TH-positive striatal subregions (i.e., lateral, central, medial, as indicated in diagram) in AAV-TYR-injected rats at 4 m post-AAV injections, either sham-treated (UT) or treated with tFUS. \**p*≤0.05 vs respective UT animals. Two-way ANOVA with Tukey’s post-hoc test. Values are mean ± SEM. N=6 animals per experimental group.

## Notes

### Competing Interest Statement

The authors have declared no competing interest.

